# Combination epigenetic-targeted therapy increases the immunogenicity of poorly immunogenic sarcomas

**DOI:** 10.64898/2026.06.18.733244

**Authors:** Alice Recho, Himavanth R. Gatla, Erin E. Resch, Maggie J. Phillips, Stephanie Glavaris, Michele Doucet, Amanda N.M. Looi, Michael I. Barbato, Nicolas J. Llosa, Michael A. Koldobskiy, Brian H. Ladle

**Author notes:** **Corresponding author:** Brian H. Ladle, CRB1 Rm 208, 1650 Orleans St., Baltimore, MD 21287, USA. Co-first authors – AR and HRG.

## Abstract

Immunotherapy approaches have shown limited efficacy in pediatric sarcomas, partly because these tumors have low mutation burden and few neoantigens. We sought to increase the immunogenicity of low mutation sarcomas by inducing expression of epigenetically silenced genes using the hypomethylating agent decitabine and histone deacetylase inhibitor entinostat. Using a mutated Kras-driven murine sarcoma model KP Sarc, sequential treatment with decitabine and entinostat significantly increased expression of silenced genes, including cancer testis antigens, and enhanced antigen presentation, including MHC I expression, compared with either agent alone. Vaccination with irradiated, epigenetically treated KP Sarc cells in a GM-CSF-secreting whole-cell vaccine induced T cell immunity against a matched tumor challenge. The anti-tumor response was directed toward epigenetically upregulated antigens, was T cell dependent, was further potentiated by immune checkpoint inhibition, and conferred immunologic memory. We showed that epigenetically regulated antigens can be shared between tumors providing protective immunity against both epigenetically treated KP Sarc and a second murine sarcoma M-3-9M. Treatment of human sarcoma lines with decitabine and entinostat induced similar gene expression changes, including shared antigen targets, and increased MHC I expression. These findings demonstrate that epigenetically upregulated antigens can serve as effective tumor-specific targets and broaden immunotherapy strategies for low-mutation sarcomas.

## Introduction

Sarcomas account for less than 1% of adult cancers, but nearly 20% of all pediatric cancers (1, 2). While standard treatment approaches combining chemotherapy, surgery, and radiation therapy are effective for many sarcoma patients with localized disease, these regimens are associated with significant short- and long-term side effects and few improvements have been made to these multimodal therapies in the last 30 years. For patients with recurrent and/or metastatic disease, overall prognosis remains dismal, emphasizing the need for improved therapies (3).

Immunotherapy has revolutionized treatment for several cancers, yet most sarcomas are resistant to these approaches. Clinical trials of immune checkpoint inhibitors (ICI) across multiple sarcoma subtypes, (particularly common pediatric sarcomas) including rhabdomyosarcoma, osteosarcoma, Ewing sarcoma, and soft-tissue sarcomas, have demonstrated limited benefit (4-7). The efficacy of ICIs depends on a pre-existing anti-tumor T cell repertoire which correlates with the tumor mutation burden and associated neoantigens (8, 9). Sarcomas often have a low mutation burden and limited neoantigen availability (10-13), further compounded by an immunosuppressive tumor microenvironment rich in inhibitory myeloid populations and cytokines (14, 15).

Epigenetic modulators have emerged as promising therapeutics capable of reshaping tumor immunogenicity. Hypomethylating agents (HMAs) such as azacitidine and decitabine (DAC) inhibit DNA methyltransferases, resulting in promoter demethylation and re-expression of silenced genes (16). Global hypomethylation can also activate endogenous intergenic elements, triggering a “viral mimicry” state with type I/III interferon signaling and increased antigen presentation (17, 18). Histone deacetylase inhibitors (HDACi) shift the balance toward increased histone acetylation and transcriptional activation, leading to broad changes in gene expression (19)] (20). HDACi has demonstrated efficacy and garnered FDA approval for cutaneous- and peripheral-T cell lymphoma but generally requires combination strategies in solid tumors [23-25]. In addition to direct tumor effects, reported impact of HDACi on immune related pathways includes increased interleukin-8 expression, increased antigen presentation pathways, and decreased suppressive effects of myeloid-derived suppressor cells (21-23).

Combining HMAs with HDACi has long been recognized to synergistically induce expression of epigenetically silenced genes, including cancer testis antigens (CTAs) (24, 25). CTAs derive from developmental proteins that are normally repressed in healthy tissues but are aberrantly expressed in some cancers such as synovial sarcoma or non-small cell lung cancer (26, 27). These antigens can serve as immune targets using a vaccine approach or engineered T cells (28, 29). While most cancers do not express CTAs at a baseline, prior work by our group and others have shown that treatment with low-dose HMAs can induce expression of CTAs in various cancer types (24, 30-32).

In this study, we hypothesize that combined HMA and HDACi treatment can increase the immunogenicity of low–mutation burden sarcomas by inducing expression of epigenetically regulated proteins capable of serving as potent T cell antigens. Using the undifferentiated murine sarcoma model KP Sarc, we show that HMA followed by HDACi treatment induces widespread methylation and transcriptional changes, leading to upregulation of antigenic proteins, enhanced MHC I expression, and activation of antigen processing pathways rendering tumor cells more susceptible to T cell killing. Similar transcriptional changes were observed in multiple human sarcoma cell lines. To test whether these induced antigens could drive immune protection, we employed the whole-cell, GM-CSF–secreting cancer vaccine (GVAX) to prime mice before challenge with epigenetically treated KP Sarc. We demonstrate that epigenetic priming combined with GVAX protects against tumor challenge, promotes tumor regression in a T cell–dependent manner when combined with ICI, and establishes long-term immune memory to epigenetically regulated antigens, including shared antigens between KP Sarc and the unrelated murine sarcoma M3-9-M.

## Results

### Sequential epigenetic therapy synergistically induces broad transcriptional activation of previously silenced genes and immune pathways *in vitro*

In these studies, we focused on the HMA DAC because of its wide-spread clinical use and the potential use of second generation HMAs under development with improved drug kinetic properties (e.g. guadecitabine). We chose the HDACi entinostat (Enti), an inhibitor of HDAC1 and HDAC3, as both of these HDAC are robustly expressed in our tumor cell line and its known immune modulatory benefits studied in other solid tumor models (21, 33-35). We aimed to determine the lowest effective concentration of Enti necessary to induce increased acetylation of histones. DAC was used at 100 nM, a dose determined to carry little toxicity while leading to hypomethylation and expression of methylation-silenced genes [35]. KP Sarc cells were treated in vitro for 72 h with increasing concentrations of Enti and assessed by Western blot for the effects on histone H3 (H3) and acetylated histone H3 (Ac-H3) and light microscopy for changes to cell morphology (**Supplemental Figure 1A-D**). 2 µM was the lowest concentration of Enti showing a significant increase in Ac-H3/Total H3 (p<0.01). Pre-treating with DAC did not further increase Enti-induced histone acetylation or change the lowest effective concentration of Enti. To verify the effect of DAC on KP Sarc, we used whole-genome bisulfite sequencing to identify the DNA mean methylation levels (MML) after control (CTL -untreated), DAC, and/or Enti in vitro treatment (**Supplemental Figure 1E**). DAC treatment led to a pronounced reduction in global methylation compared to untreated or Enti-treated cells. This hypomethylation was prominent near transcription start sites (TSSs), where CpG islands are typically enriched. Enti alone had minimal effect on DNA methylation, while sequential DAC then Enti treatment (DAC+Enti) exhibited moderate hypomethylation.

RNA sequencing of treated cells identified patterns of differentially expressed genes (DEGs) following each treatment. Differential expression analysis using a cutoff of 2.5 log₂ fold change and adjusted p value (padj) < 0.05 revealed the expected pattern of many more upregulated genes compared to downregulated genes in treated KP Sarc cells (**Figure 1A**). Notably, DAC+Enti induced the largest number of upregulated genes, many of which were initially expressed at a low level in untreated KP Sarc, suggesting strong transcriptional activation of previously silenced genes. Among these upregulated genes were canonical CTAs (**Supplemental Figure 1F**) with the most robust expression present in the DAC+Enti treated cells. 590 of the DAC+Enti upregulated genes were unique to the combination treatment (**Figure 1B**), with substantial overlap of 551 genes with Enti-treated cells, suggesting that Enti contributes more to the upregulated gene expression in the DAC+Enti treated cells with DAC treatment resulting in many genes poised for expression. With our focus on the potential antigenicity of newly induced genes, we analyzed genes with minimal expression in control KP Sarc cells which are upregulated in one of the treatment groups (**Figure 1C, Supplemental Table 1**). Hierarchical clustering grouped the genes into 4 main groups consistent with differing mechanisms of gene expression regulation. Group 1 contained genes where DAC alone was sufficient to induce expression. Of note, all but 3 of these genes further induced expression upon subsequent Enti treatment. The expression of Group 2 genes was driven entirely by Enti exposure. Group 3 included genes which required both DAC and Enti for expression – potentially requiring promoter demethylation resulting in a gene poised for expression upon HDACi treatment. Group 4 consisted of genes whose expression was mildly induced by Enti alone, but pretreatment with DAC allowed for much more robust expression with Enti exposure. We investigated gene sets enrichment of GO Biological Processes (**Figure 1D**) induced in DAC+Enti treated tumors that were shared by DAC or Enti alone-treated tumors. While Enti alone shared enriched pathways like “export across plasma membrane”, the DAC+Enti combination uniquely enriched immune-related pathways such as “antigen processing and presentation of peptide antigen” and “inflammatory response to antigenic stimulus.” Furthermore, DAC and DAC+Enti shared enrichment of viral defense pathways like “defense response to virus,” suggesting a DAC-driven viral mimicry signature (17). These data suggest that DAC+Enti treatment of KP Sarc tumor cells leads to more robust transcriptional activation, particularly in immune and antigen presentation pathways that may portend enhanced anti-tumor immunity through the epigenetically induced pathways and neoantigen expression.

**Figure 1:**
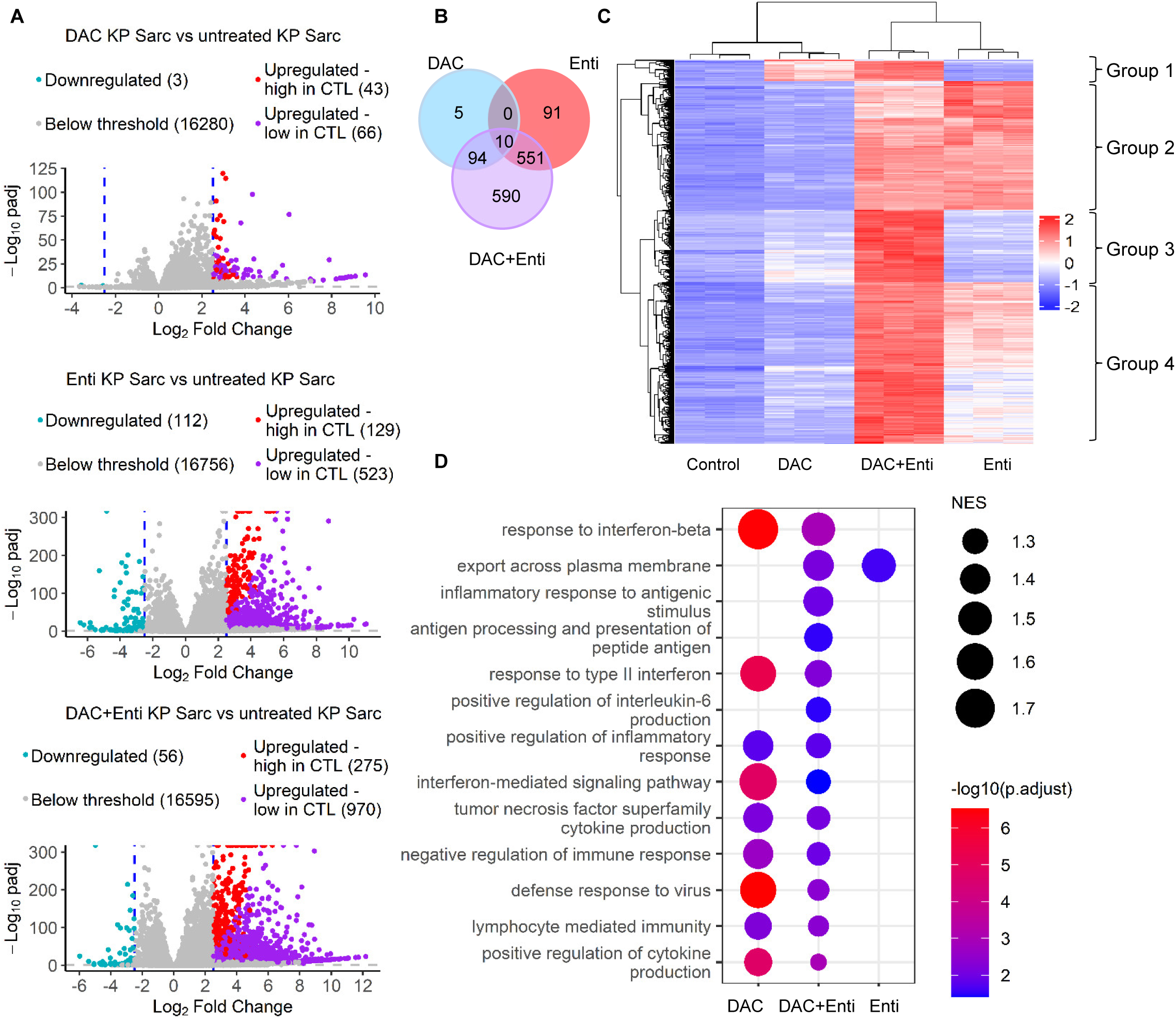
Sequential epigenetic therapy induces broad gene expression changes in KP Sarc cells. *(A)* Volcano plots of DAC, Enti and DAC+Enti treated KP Sarc compared to CTL KP Sarc showing the number of differentially expressed genes that met thresholds of -2.5 and 2.5 log2 fold change and padj<0.05 after normalization. “Upregulated -high in CTL” genes are defined as genes with normalized average expression counts in CTL tumor cells above 50. “Upregulated -low in CTL” genes are defined as genes with normalized average expression counts below 50 in CTL and above 50 in DAC+Enti tumor cells. *(B)* Venn diagram of all upregulated genes in each of DAC, Enti and DAC+Enti treated KP Sarc, as visualized on the volcano plots in (A). *(C)* Heatmap of the upregulated, low in CTL genes identified in (A). Count data were normalized and scaled using Z-score. *(D)* Bubble plot of significantly enriched GO Biological Process pathways from GSEA comparing each of DAC, Enti and DAC+Enti treated KP Sarc to CTL. Bubble size indicates normalized enrichment score (NES) and color scale indicates p adj value.

### *In vitro* treatment alone does not render KP Sarc tumors more immunogenic upon *in vivo* tumor challenge

With the significant gene expression changes noted above – including potential increased tumor antigen processing – we explored whether the in vitro treated KP Sarc tumors would have decreased fitness or increased baseline anti-tumor immune activation in vivo. We injected an equal dose of in vitro-treated KP Sarc tumor cells into the gastrocnemius as an orthotopic site of tumor challenge in BL/6 mice and measured leg circumference every two days. There was no significant difference between the average tumor growth of DAC-treated, Enti-treated, or DAC+Enti-treated KP Sarc tumors compared to untreated KP Sarc tumors (**Figure 2A**). Moreover, we characterized the tumor infiltrating immune cells by flow cytometry once tumors reached maximal size. We found no significant differences among the four treatment conditions in the numbers of total CD45^+^ hematopoietic-derived cells, nor the composition of those immune cells with similar proportions of myeloid CD11b^+^ cells and T cells (CD8^+^, CD4^+^Foxp3^-^ and CD4^+^Foxp3^+^) (**Figure 2B**). Thus, the gene expression changes induced in vitro are not associated with increased immune activation upon simple exposure to the orthotopically injected treated tumor cells. Importantly, the in vitro treatment did not significantly alter the tumor cell fitness, as equal rates of tumor growth were observed.

**Figure 2:**
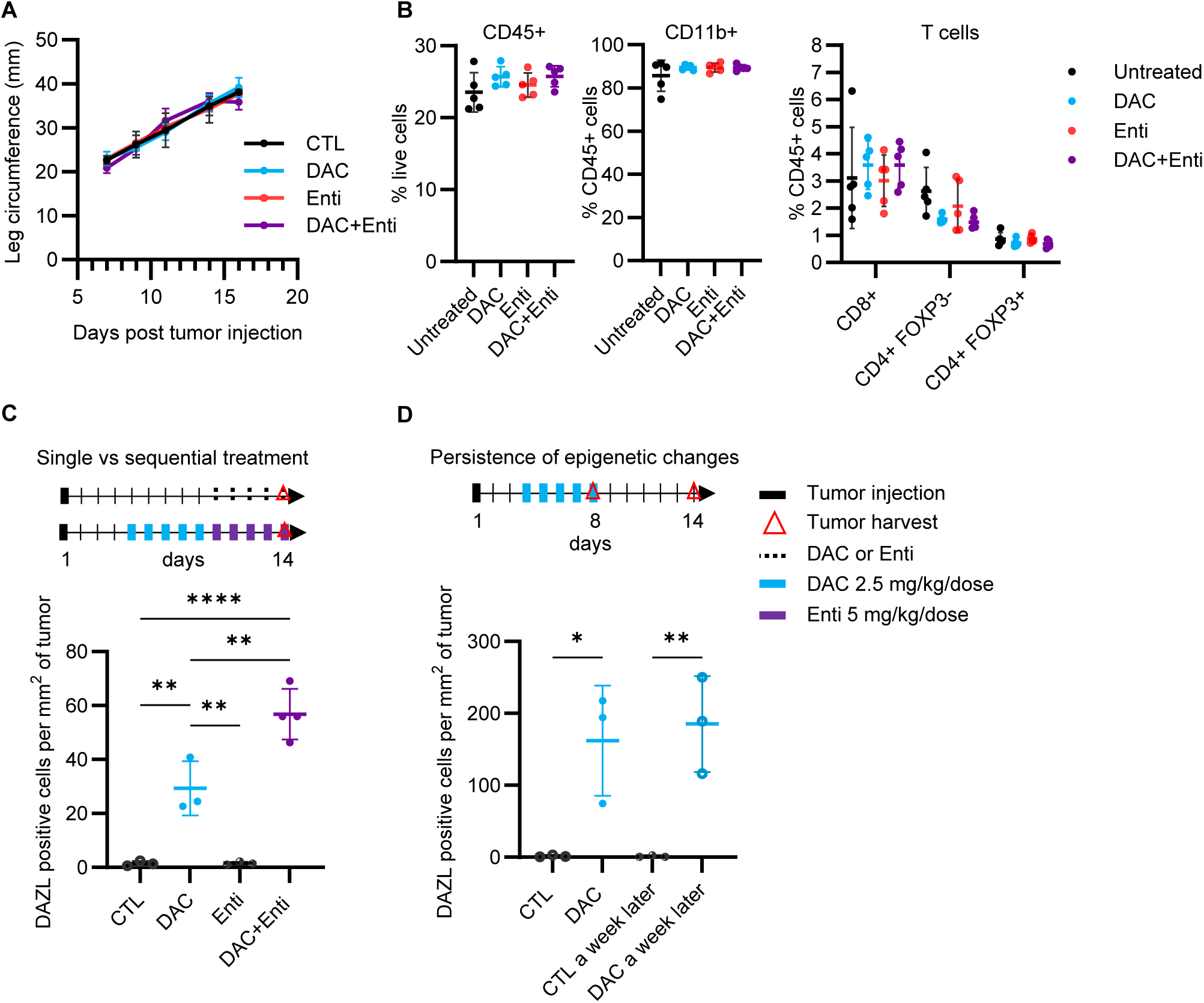
Epigenetic treatment of KP Sarc does not impact tumor fitness or tumor microenvironment. *(A*) Average leg circumference of BL/6 injected with KP Sarc treated in vitro with DAC, Enti, or DAC+Enti. There is no significant difference between the curves as determined by two-way ANOVA with Tukey’s multiple comparison test. *(B)* Quantification using flow cytometry of CD45^+^, CD11b^+^ cells and T cells of KP Sarc harvested when maximal size was reached. No significant differences were observed as determined by one-way ANOVA with Tukey’s multiple comparison test. (C) Experimental schema of in vivo single or sequential treatment of mice bearing KP Sarc; and expression density analysis results from IHC staining for Dazl positive cells per mm^2^ of analyzed tumor. (D) Experimental schema of in vivo single-agent DAC treatment of mice bearing KP Sarc; and expression density analysis results from IHC staining for Dazl positive cells per mm^2^ of analyzed tumor. Tumors were harvested at the completion of the five days of DAC treatment or six days later. *p<0.05, **p<0.01, ***p<0.001, ****p<0.0001 as determined by one-way ANOVA with Tukey’s multiple comparison test.

We next asked whether the epigenetic changes observed in KP Sarc tumors with in vitro HMA and HDACi exposure could also be induced and sustained in vivo. We treated KP Sarc-bearing mice with either DAC, Enti or sequential DAC+Enti for five days before harvesting tumors at day 14 when maximum tumor diameter of 2 cm was reached (**Figure 2C**). Harvested tumors were evaluated for expression of representative murine CTAs, Dazl and Trap1a for which we knew adequate hypomethylation would be required for expression (**Supplemental Figure 1F**). There was significant upregulation of *Dazl* and *Trap1a* RNA expression after DAC and DAC+Enti treatment in vivo as assessed by quantitative PCR (**Supplemental Figure 2A**). Correlating with induction of *Dazl* transcription, a significant increase in Dazl protein expression was confirmed via IHC of tumor tissue after in vivo DAC treatment, with higher expression in DAC+Enti treated tumors, compared to no expression in CTL or Enti cohorts (**Figure 2C**). To confirm if DAC-induced expression changes persist, we examined *Dazl* and *Trap1a* gene expression by qPCR **(Supplemental Figure 2B**) and Dazl protein expression by IHC (**Figure 2D**) of in vivo treated tumors at the end of treatment and one week later. Since KP Sarc tumors reach a maximal size by day 14 and sequential DAC+Enti treatment takes ten days, we could only test DAC-alone treated cohorts. IHC and qPCR confirmed persistent expression of these DAC-induced genes even one week after completing treatment.

### Active immune priming with DAC-, Enti-, and DAC+Enti-treated GVAX decreases the growth of similarly treated KP Sarc tumors further potentiated by immune checkpoint inhibitor therapy

We have established that our epigenetic therapy can upregulate newly expressed antigens, but in vivo exposure to the epigenetically modified tumors is not sufficient to generate anti-tumor immunity. We hypothesized that a potent immune response can be induced against antigens upregulated by epigenetic therapy using a vaccine approach. Using our irradiated whole-cell tumor vaccine approach, we made four vaccine cocktails combining our GM-CSF-secreting bystander cell line with: 1. control KP Sarc cells (Untreated KP Sarc GVAX), 2. DAC-treated KP Sarc (DAC KP Sarc GVAX), 3. Enti-treated KP Sarc (Enti KP Sarc GVAX) or 4. DAC+Enti-treated KP Sarc (DAC+Enti KP Sarc GVAX). Tumor challenge occurred seven days after GVAX administration to allow for immune activation (**Figure 3A**). To determine if GVAX-induced immune responses were specific to epigenetically regulated antigens, each mouse was challenged orthotopically in the gastrocnemius with untreated KP Sarc cells or KP Sarc cells pre-treated in vitro with the same condition (DAC alone, Enti alone, or DAC+Enti) as the GVAX used for priming. Tumor growth was measured every two days. As we know GVAX induces T cell responses (36), we hypothesized that anti-tumor activity would be further augmented by ICI. Thus, an additional cohort of GVAX-treated mice started anti-PD1 and anti-CTLA-4 treatment the day after tumor challenge.

**Figure 3:**
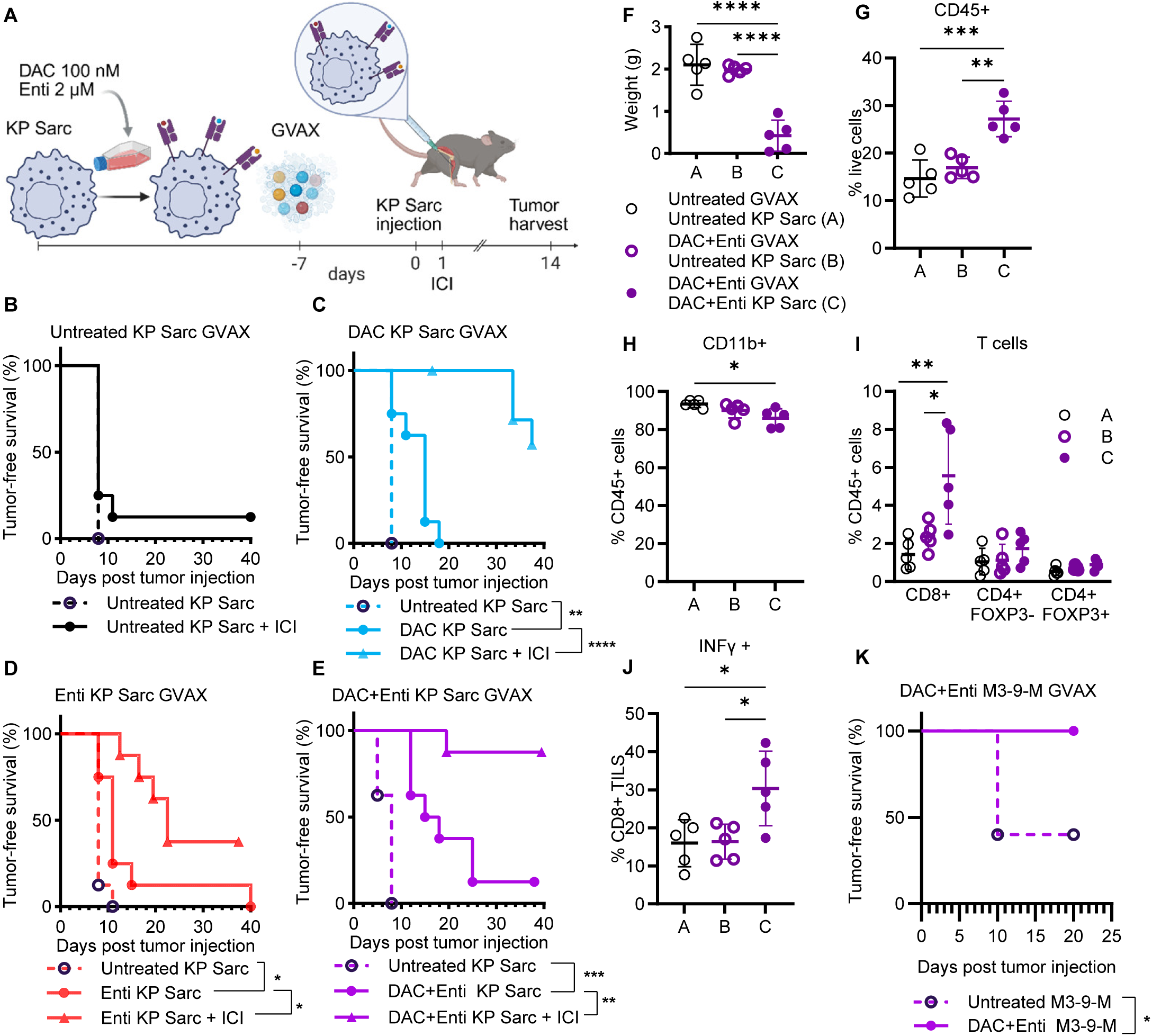
Priming mice using epigenetically-treated GVAX decreases tumor growth induced by similarly treated KP Sarc cells and further potentiated by immune checkpoint inhibition (ICI). *(A)* Experiment schema showing GVAX vaccination of BL/6 mice (n=8), one week before intramuscular injection with similarly treated KP Sarc, followed by ICI treatment. *(B)* Kaplan-Meier curves showing tumor-free survival of BL/6 mice which were primed with untreated KP Sarc GVAX and were challenged with untreated KP Sarc with or without ICI. (C) Kaplan-Meier curves showing tumor-free survival of BL/6 mice which were primed with DAC GVAX and were challenged with untreated KP Sarc or DAC treated KP Sarc with or without ICI. *(D)* Kaplan-Meier curves showing tumor-free survival of BL/6 mice which were primed with Enti GVAX and were challenged with untreated KP Sarc or Enti treated KP Sarc with or without ICI. *(E)* Kaplan-Meier curves showing tumor-free survival of BL/6 mice which were primed with DAC+Enti GVAX and were challenged with untreated KP Sarc or DAC+Enti treated KP Sarc with or without ICI. *(F)* Weight of untreated KP Sarc tumors harvested after untreated GVAX (A) or DAC+Enti GVAX (B), and of DAC+Enti KP Sarc harvested after DAC+Enti GVAX (C). Figure legend applies for figures G-J. Quantification using flow cytometry of CD45^+^ (*G*) as percentage of live cells, CD11b+ (H), CD8^+^, CD4^+^ FOXP3^-^ and CD4^+^ FOXP3^+^ T cells (I) and IFN-γ^+^ percent of CD8^+^ TILs (J). The values represent the mean +/- SD. *p<0.05 as determined by one-way ANOVA with Tukey’s multiple comparison test. (K) Kaplan-Meier curves showing tumor-free survival of BL/6 mice primed with M3-9-M GVAX and injected with DAC+Enti treated M3-9-M. The values represent the mean +/- SD. *p<0.05, **p<0.01, ***p<0.001 as determined by two-sided log-rank test.

Untreated KP Sarc GVAX provided no protection against untreated KP Sarc tumor challenge and was not improved with ICI treatment. These findings suggest the poor immunogenicity of the untreated KP Sarc tumors (**Figure 3B**). After DAC GVAX, mice that received DAC pre-treated KP Sarc had a significantly improved tumor-free survival as compared to those challenged with untreated KP Sarc (**Figure 3C**) but all mice succumbed to tumor. Survival was significantly improved by adding ICI with 60% of mice never developing tumor. Similarly, after Enti GVAX, mice had significant delay in developing Enti-treated KP Sarc tumors with the addition of ICI resulting in complete rejection of the Enti-treated tumors in 35% of mice (**Figure 3D**). We observed the most robust protection from GVAX in the DAC+Enti GVAX-treated mice. As in all other conditions, the DAC+Enti GVAX-treatment provided no protection upon challenge with untreated KP Sarc tumors. Yet, the DAC+Enti GVAX resulted in a significant delay in DAC+Enti-treated KP Sarc tumors with ICI boosting the response further with 85% of mice completely rejecting the DAC+Enti-treated KP Sarc tumors (**Figure 3E** and **Supplemental Figure 3**). These findings provide strong evidence for vaccine-induced immune responses to epigenetically regulated antigens capable of providing potent anti-tumor immunity. Separate cohorts of mice were treated as above and harvested day 14 of tumor growth to evaluate tumor infiltrating immune cell populations. Consistent with improved tumor-free survival in the DAC+Enti GVAX-treated mice, we observed significantly lower tumor weights in the DAC+Enti-treated KP Sarc tumors compared to untreated GVAX or untreated KP Sarc GVAX tumors (**Figure 3F**). DAC+Enti GVAX resulted in increased infiltration of CD45^+^ immune cells (**Figure 3G**), a decreased proportion of CD11b^+^ myeloid cells (**Figure 3H**) and increased proportions CD8^+^ T cells (**Figure 3I**) only in the DAC+Enti tumors compared to control KP Sarc tumors or control GVAX-treated mice. While no significant differences were observed in immune checkpoint expression on the tumor infiltrating CD8^+^ T cells (**Supplemental Figure 4A** and **4B**), we did find a significant increase in the effector IFN-γ-producing CD8^+^ T cells in the DAC+Enti tumors (**Figure 3J**).

We validated these findings using a second syngeneic murine sarcoma model M3-9-M (37). BL/6 mice were vaccinated with DAC+Enti-treated M3-9-M GVAX. One week later, mice were challenged with untreated M3-9-M or DAC+Enti-treated M3-9-M and followed for tumor growth. Consistent with the known immunogenicity of M3-9-M (37), DAC+Enti GVAX provided some protection against untreated M3-9-M with 40% of mice remaining tumor-free. However, the 100% protection against DAC+Enti-treated M3-9-M tumor cells provides further evidence that DAC+Enti GVAX induced potent immune responses against epigenetically regulated antigens (**Figure 3K**).

### DAC+Enti-induced tumor associated antigens (TAAs) can elicit long-term adaptive anti-tumor immunity

To investigate if epigenetically induced tumor-associated antigens are capable of generating long-term memory anti-tumor immune responses, we re-challenged the mice which rejected tumors in the experiments described above. After a minimum of 60 days, mice that previously rejected tumors were re-challenged with both untreated (one leg) and DAC+Enti treated (opposite leg) KP Sarc. Control KP Sarc tumors grew while 100% of the mice rejected DAC+Enti KP Sarc in the other leg (**Figure 4A**). Similar results were observed in mice that have rejected DAC+Enti M3-9-M cells and were re-challenged with untreated and DAC+Enti-treated M3-9-M (**Figure 4B**) indicating an immune memory to epigenetically upregulated antigens. We hypothesized that epigenetically regulated tumor antigens could be shared between unrelated syngeneic tumor models. To investigate if this immune memory can be established against shared antigens, we rechallenged DAC+Enti KP Sarc GVAX-treated mice that had previously rejected DAC+Enti KP Sarc with untreated and DAC+Enti treated M3-9-M cells (**Figure 4C**). Prior DAC+Enti KP Sarc GVAX prevented growth of DAC+Enti M3-9-M while untreated M3-9-M grew normally indicating immune memory to shared epigenetically regulated antigens capable of being induced in both KP Sarc and M3-9-M tumors. Our experiments showing improved vaccine responses with ICI and immune memory suggest a T cell-mediated anti-tumor immunity. To further explore the role of T cells in preventing tumor growth, we depleted CD4^+^ and/or CD8^+^ T cells one week after DAC+Enti GVAX vaccination (**Supplemental Figure 4C** and **4D**). Depleting CD4^+^ and CD8^+^ T cells ablates any benefit in the DAC+Enti GVAX-treated mice (**Figure 4D**) showing that the tumor-specific immune response to tumor-associated antigens is T cell mediated.

**Figure 4:**
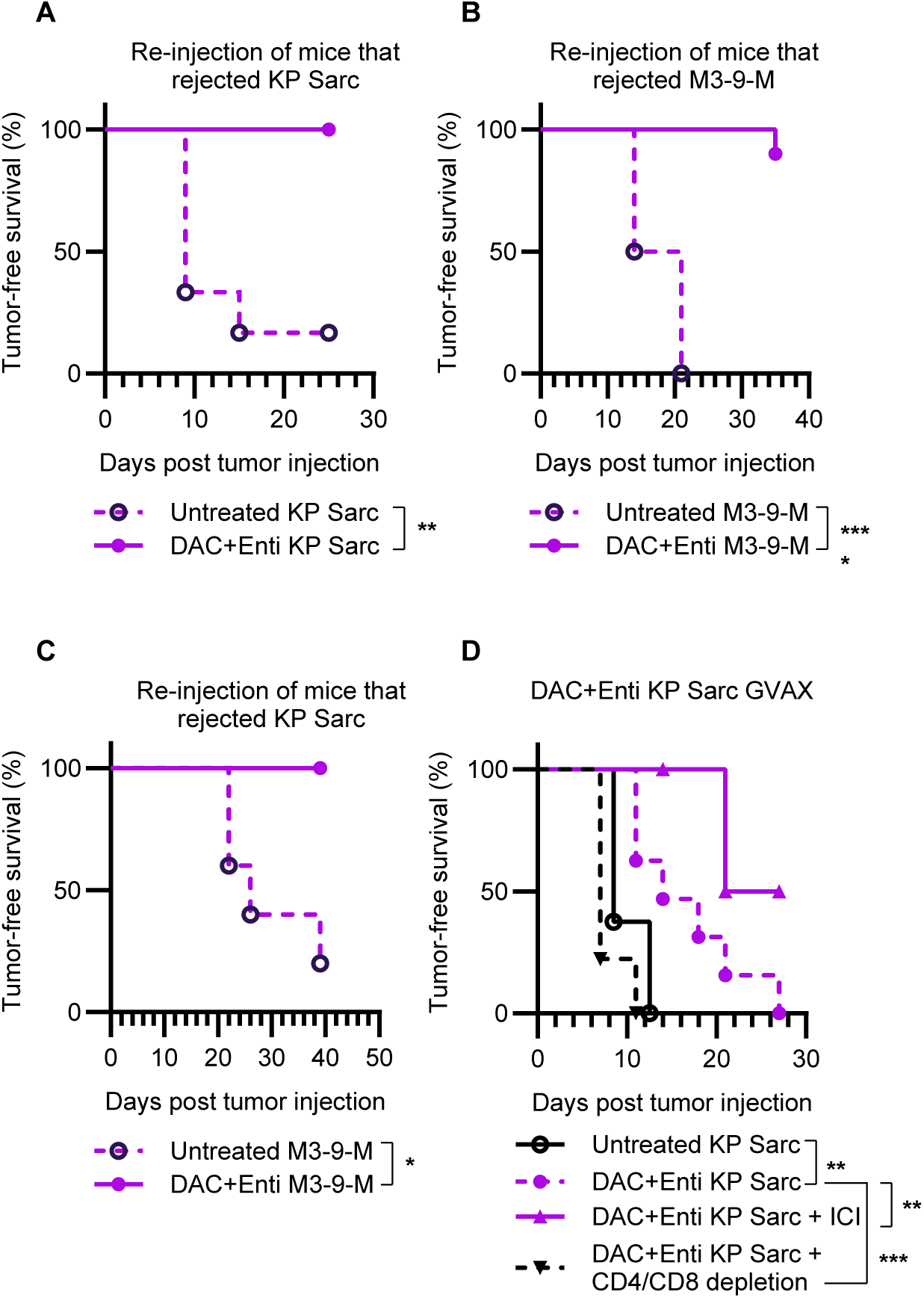
Epigenetically-induced tumor associated antigens elicit a long-term adaptive anti-tumor immune response. *(A)* Kaplan-Meier curves showing tumor-free survival of BL/6 mice which were previously primed with DAC+Enti GVAX and rejected KP Sarc tumor growth and subsequently injected with a rechallenge of untreated or DAC+Enti treated KP Sarc. *(B)* Kaplan-Meier curves showing tumor-free survival of BL/6 mice which were previously primed with DAC+Enti GVAX and rejected M3-9-M tumor growth and subsequently injected with a rechallenge of untreated or DAC+Enti treated M3-9-M. *(C)* Kaplan-Meier curves showing tumor-free survival of BL/6 mice which were previously primed with DAC+Enti GVAX and rejected KP Sarc tumor growth and subsequently injected with a rechallenge of untreated or DAC+Enti treated M3-9-M. (*D)* Kaplan-Meier curves showing tumor-free survival of BL/6 mice which were primed with DAC+Enti GVAX and were injected with DAC+Enti treated KP Sarc with or without ICI, or received CD4^+^ and CD8^+^ depleting antibodies. The values represent the mean +/- SD. *p<0.05, **p<0.01, ***p<0.001 as determined by two-sided log-rank test.

### HDACi-induced MHC-I expression is not sufficient to increase immunogenicity of treated KP Sarc

As described in **Figure 1**, broad gene expression changes occur with DAC+Enti treatment. In addition to induction of potential novel antigens, pathways of antigen processing/presentation are also upregulated, likely contributing to the immunogenicity of the treated tumors. We analyzed the effects of epigenetic therapy specifically on surface MHC-I expression on the tumor cells known to be a key requirement for T cell recognition. While DAC treatment mildly increases MHC-I expression, Enti increases the MFI of MHC-I expression more than 7-fold compared to untreated KP Sarc and DAC+Enti treatment increases more than 16-fold (**Figure 5A**). To explore if increased MHC-I expression was sufficient to explain the increased immunogenicity of the DAC+Enti-treated tumors, we utilized another pathway known to induce MHC-I expression with in vitro IFN-γ treatment which increased MHC-I expression even beyond that seen with Enti- or DAC+Enti-treatment (**Figure 5B**). Mice vaccinated with DAC+Enti KP Sarc GVAX were then challenged with untreated KP Sarc, IFN-γ-treated KP Sarc, or DAC+Enti-treated KP Sarc. Vaccinated mice were still unable to control tumor growth of IFN-γ-treated KP Sarc despite high levels of MHC-I (**Figure 5C**). Thus, while the increase of antigen presentation pathways likely improves immune recognition of DAC+Enti treated tumors, expressed epigenetically regulated antigens are the targets of the vaccine-induced T cell response.

**Figure 5:**
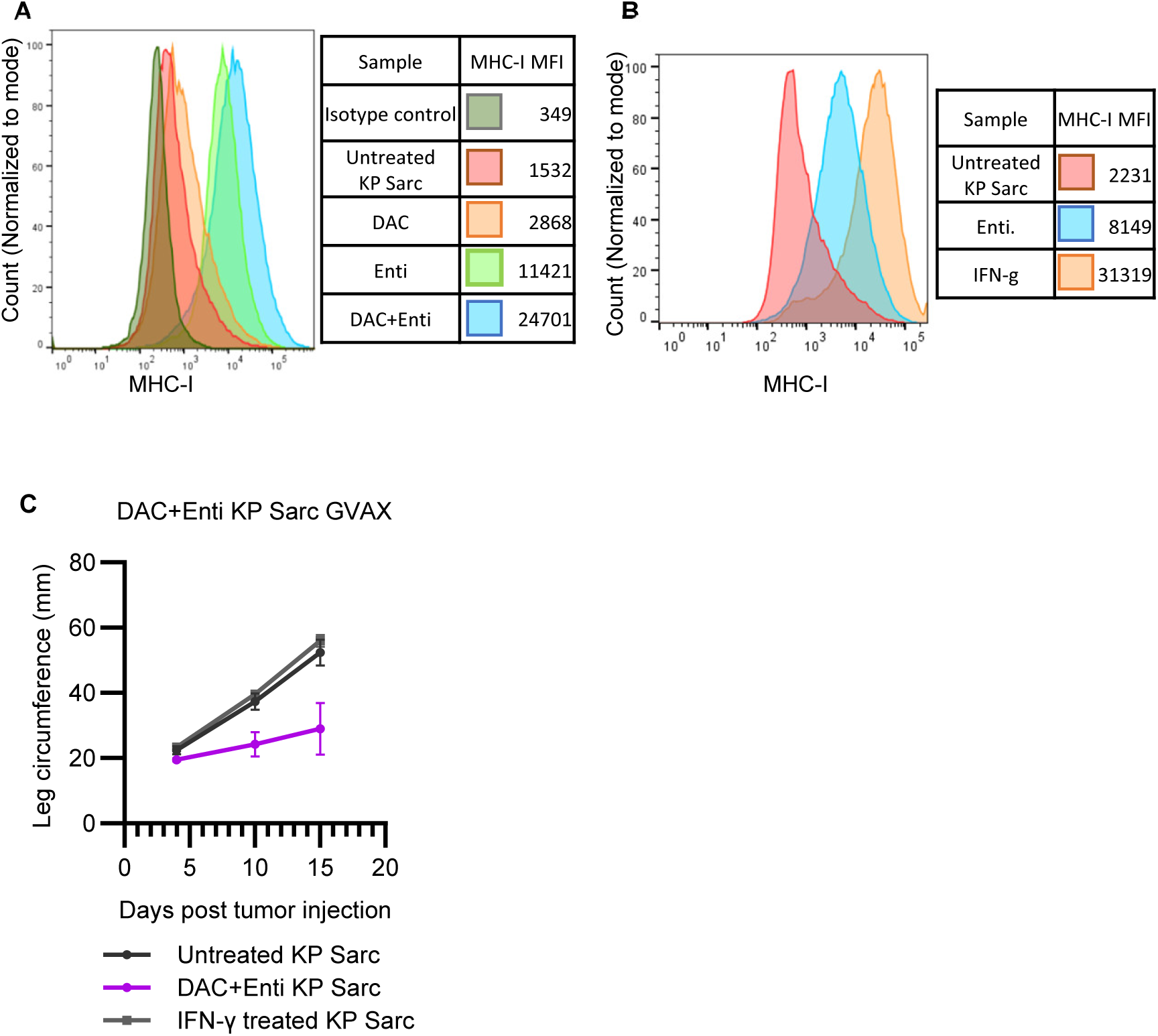
HDAC inhibition induces the expression of MHC-I. *(A)* Expression of MHC-I in KP Sarc cells treated in vitro with DAC and/or Enti and quantified using flow cytometry. (*B)* Expression of MHC-I in KP Sarc cells treated in vitro with Enti or with IFN-γ for 24 h, and quantified using flow cytometry. *(C)* Average leg circumference of BL/6 mice injected with untreated, DAC+Enti treated or IFN-γ treated KP Sarc, with pairwise comparisons determined by nonlinear regression.

Shared antigens are upregulated in response to DAC+Enti in human Ewing sarcoma and rhabdomyosarcoma models.

To explore the translational relevance and uncover potential immune targets in human sarcomas, we extended our analysis to DAC+Enti treated *PAX3::FOXO1* fusion-positive rhabdomyosarcoma cell lines RH30 and RH41, and *EWSR1::FLI1* expressing Ewing sarcoma cell line CHLA-9. Bulk RNA sequencing revealed a pronounced skew towards upregulated genes (**Figure 6A**) with many of these upregulated genes expressed at low levels in control conditions, consistent with our murine KP Sarc cell line. Principal component analysis (PCA) of significantly differentially expressed genes demonstrated clear separation between treated and untreated samples (**Figure 6B)** indicating substantial transcriptional shifts associated with treatment across cell lines. As noted in the murine KP Sarc tumor cells, DAC+Enti increased mechanisms of antigen presentation such as increased *HLA-A, -B*, and *-C* with particular induction of HLA genes poorly expressed in the untreated cells (e.g. *HLA-B* in RH41 and CHLA-9, and *HLA-C* in RH41) (**Figure 6C**). Exploring gene expression changes that may represent potential tumor-restricted antigens, we analyzed expression of a curated list of CTAs (38) in untreated versus DAC+Enti treated tumor cells (**Supplemental Figure 5, Supplemental Table 2**). Some CTAs were expressed at high levels in untreated tumor cells with little change upon DAC+Enti treatment (e.g. *CEP55, KIF2C, PBK*). Other CTAs were only upregulated in one or two of the tumor cell lines, but not shared between all three (e.g. *DAZL, GAGE2A, FMR1NB*). The well-known and clinically targeted CTA *PRAME* was expressed at high levels at baseline in the EWS line, CHLA-9, and DAC+Enti induced high expression levels in RH30 and RH41 making it a viable shared CTA. A notable block of CTAs met criteria of being significantly upregulated in all three tumor cell lines (**Figure 6A**) and expressed at very low levels in untreated tumors (see **Supplemental Table 3** for normalized log_2_(TPM+1) expression values) including *MAGEA4* (the target of FDA-approved adoptive T cell therapy afamitresgene autoleucel), *BRDT*, *CTAG2*, *CTFCL*, *GPAT2*, *MAEL*, and other *MAGE* isoforms. To further assess overlap in treatment-induced gene expression changes beyond CTAs, a Venn diagram of upregulated genes across RH30, RH41, and CHLA-9 showed a set of 240 genes upregulated in all three cell lines (**Figure 6D, Supplemental Table 3**). While the antigenic potential of some of these upregulated genes is low (e.g. T cell coreceptors *CD4* and *CD8A* whose expression is known to be controlled by DNA methylation (39)), we hypothesize that potential epigenetically-regulated tumor-restricted T cell targets can extend beyond CTAs. One example includes *LRRC15*, encoding a tumor-restricted, surface-expressed, protein that is significantly increased or induced in these DAC+Enti treated sarcoma cell lines that is being studied as a target of adoptive cell therapies (40), antibody targeting approaches (41), but also plays a key role in the immune modulation of the tumor microenvironment (42, 43). It is important to consider that some gene expression changes induced by DAC+Enti may be detrimental. Examples include *INHBA*, which contributes to metastatic spread and TGF-beta signaling in breast cancer (44) and is a poor prognostic indicator in several cancers (45, 46), and the lncRNA *LINC00624*, shown to cause immunosuppressive changes in HER2+ breast cancer models (47). While exploratory, these findings highlight shared transcriptional changes activated by DAC+Enti treatment, suggesting common molecular responses to epigenetic modulation in human sarcomas. This data indicates the potential for common shared antigens that could be immunologically targeted between individual patients and different diseases as well as changes in tumor biology that need to be further characterized and understood.

**Figure 6:**
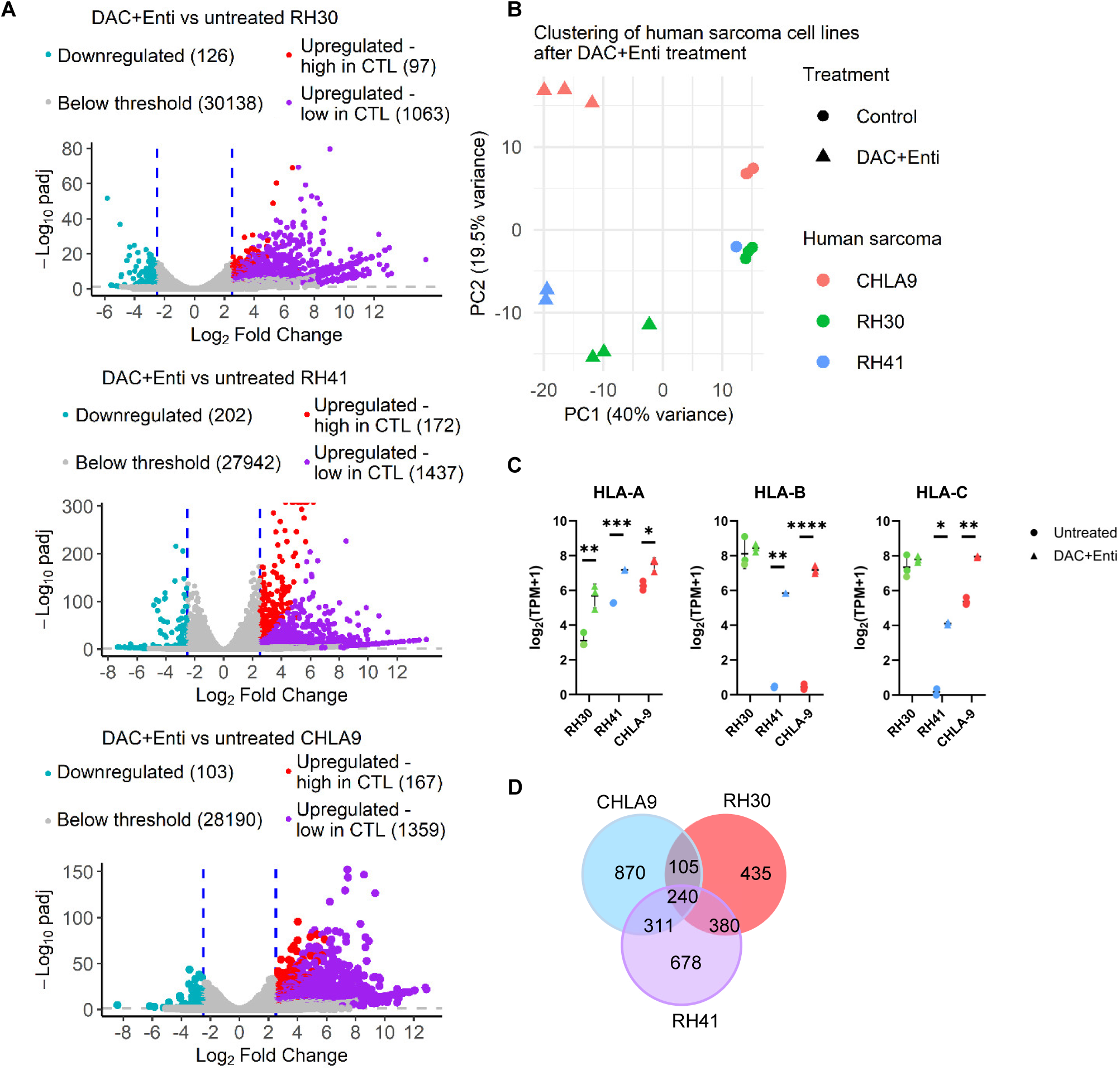
Shared antigens are upregulated in response to DAC+Enti in human Ewing sarcoma and rhabdomyosarcoma cell lines. *(A)* Volcano plots of each of DAC+Enti treated RH30, RH41, and CHLA-9 compared to untreated CTL, showing differentially expressed genes that met thresholds as described in Figure 1. *(B)* Principal component analysis (PCA) of DAC+Enti treated RH30, RH41 and CHLA-9. Only genes with padj < 0.05 and common across all three cell lines were included. Each point represents a biological replicate, colored by cell line and shaped by treatment. Axes show the first two principal components with the percentage of variance. *(C) HLA-A, -B*, and *-C* gene expression changes induced in each cell line by DAC+Enti treatment with counts normalized to log_2_(TPM+1). *p<0.05, **p<0.01, ***p<0.001, ****p<0.0001 as determined by one-way ANOVA with Tukey’s multiple comparison test. *(D)* Venn diagram of upregulated genes in each of DAC+Enti treated RH30, RH41, and CHLA-9 cells as analyzed by DESeq2 and visualized on the volcano plots in *(A)*.

## Discussion

Current treatment of sarcomas remains largely limited to cytotoxic chemotherapy, surgery, and radiation, with no improvement in survival outcomes over recent decades. While effective immunotherapy options for these cancers have remained elusive, our understanding of the obstacles that must be overcome continues to increase. Our work investigates one approach to a major limitation in sarcoma immunotherapy – the inherent poor immunogenicity of pediatric sarcomas. A common feature of cancers for which immunotherapies such as ICI have been effective is a high tumor mutation burden (9) with examples including melanoma, non-small cell lung cancer, and MSI-high colorectal cancer (27, 48, 49). Tumor mutations can result in the expression of mutated, cancer-specific proteins from which cancer-specific epitopes can be presented to the immune system. This can generate an anti-cancer immune response, leading to an inflamed tumor microenvironment, activation of adaptive immunity, and, in some cases, tertiary lymphoid structure formation, which correlates with therapeutic benefit from ICI. This trend holds true in the expansive list of sarcoma diagnoses. Studies have identified sarcomas with increased immune cell enrichment, including the presence of tertiary lymphoid structures (14, 50, 51), often seen in sarcoma subtypes known to have high tumor mutation burdens, such as angiosarcoma and undifferentiated pleomorphic sarcoma (UPS) (52). Immune-enriched sarcomas have shown response rates to ICI (30% overall response rate) similar to those seen in other ICI-responsive cancers (53). However, common pediatric sarcomas such as Ewing sarcoma, osteosarcoma, and rhabdomyosarcoma generally have few expressed mutated proteins (54), which likely contributes to the low T cell infiltration (14, 15, 55, 56) and lack of responses with ICI therapy (4-7).

With the specific goal of increasing the available antigen repertoire in poorly immunogenic sarcomas, our study explores the role of epigenetic-modifying drugs DAC and Enti to upregulate TAAs and increase antigen presentation to increase the immunogenicity of sarcomas. While DAC or Enti alone had an effect, sequential treatment with DAC followed by Enti produced the strongest upregulation of potential antigens, inflammatory pathways, and antigen presentation pathways. This synergistic interaction was reflected both in the number of upregulated genes and in gene set pathway enrichment, with DAC+Enti uniquely inducing immune-related pathways, including antigen processing and presentation, as well as potential antigen targets. Previous studies in multiple model systems have shown that low dose epigenetic therapy using HMA and HDACi can upregulate genes involved in various immunologically relevant pathways, including TAAs (18, 23, 30, 57, 58). CTAs are one well-characterized class of TAAs and have been targeted in immunotherapy approaches across cancers as diverse as glioblastoma (59), melanoma (60), triple-negative breast cancer (61), and lung cancer (62). Our data indicate that a combination such as DAC+Enti can induce broad gene expression changes beyond CTAs, allowing for many more potential antigen targets, including non-canonical gene products that may also contribute to neoantigen pools and immune recognition.

While changes in gene expression in these sarcomas were expected upon DAC+Enti treatment, our key finding showed that these epigenetically regulated gene products can serve as potent immune targets enabling tumor rejection. With the whole-cell GVAX vaccine platform, DAC+Enti-treated GVAX generated protective immunity that was selective for epigenetically regulated targets, as the vaccine afforded no protection when mice were challenged with untreated control tumor cells. The protective immunity in our models was T cell dependent, based on T cell depletion experiments ablating the anti-tumor response, augmentation of the response by ICI, and mice showing immune memory upon rechallenge. Our experiments showing shared immunity in DAC+Enti KP Sarc GVAX-treated mice against DAC+Enti-treated M3-9-M further support the conclusion that epigenetically regulated genes were the targets of DAC+Enti GVAX-induced immunity. Our ongoing and future work seeks to identify the specific epigenetically regulated antigens serving as T cell targets in our models. Based on the data presented here, we hypothesize that these are genes that are silenced in untreated sarcoma cells and newly upregulated after treatment. Candidates may include CTAs, but could also include non-canonical genes known to be upregulated with this treatment (17). Other mechanisms could include altered antigen processing that may occur with HMA or HDACi treatment that could shunt other proteins to be presented by MHC I that are not normally presented. Our studies showing cross-tumor immunity in murine GVAX experiments suggest the presence of shared antigens upregulated by DAC+Enti. We explored shared epigenetically regulated antigens in three human sarcoma cell lines – RH30 (*PAX3::FOXO1* fusion-positive RMS), RH41 (*PAX3::FOXO1* fusion-positive RMS), and CHLA-9 (*EWSR1::FLI1* positive Ewing sarcoma) – revealing a robust and distinct transcriptional response to DAC+Enti treatment, as well as a set of shared upregulated transcripts across all three lines. Although limited, these findings suggest the potential for immune-targeting strategies against shared antigens broadly applicable across multiple sarcoma subtypes.

Importantly, epigenetically treated KP Sarc tumor cells alone did not trigger an anti-tumor immune response when injected in vivo, with no delay in tumor growth or increase in immune cell infiltration compared with untreated KP Sarc. Simply inducing expression of new genes with HMA + HDACi in the KP Sarc sarcoma cells did not induce immune priming or T cell activation against them. Because the immune context of antigen presentation is crucial for T cell fate decisions during activation, our findings support the need for active immune priming against epigenetically regulated antigens, such as a vaccine or immune adjuvant, in clinical studies where the intended goal of HMA and/or HDACi is synergy with immunotherapy. Reviews have highlighted multiple clinical trials in solid tumors investigating the combination of epigenetic-modifying drugs and ICI or other immunotherapies, with limited success (63, 64). As our DAC+Enti KP Sarc GVAX experiments showed, maximal immune activation and, thus, tumor rejection required epigenetic modifiers, active vaccination, and ICI, whereas any single modality, or even paired approaches, was insufficient.

Beyond changes in antigen expression to alter sarcoma immunogenicity, we also found that DAC+Enti treatment enhances MHC I expression and induces gene expression changes associated with interferon responses, increased antigen processing and presentation, and other inflammatory pathways. These changes improve T cell recruitment and T cell-mediated tumor recognition. This is consistent with data showing that Trichostatin A (Class I and II HDACi) mediated induction of MHC I in melanoma (65). Previous studies have shown similar findings, with HDACi and HMA increasing type I IFN response in a poorly immunogenic ovarian cancer model (66). While these changes likely improve the responsiveness of our tumor models to T cell attack, we show that the anti-tumor response was not dependent on IFN-γ signaling, as IFN-γ pretreatment failed to reproduce tumor rejection in the absence of DAC+Enti exposure, emphasizing the essential role of epigenetically upregulated TAAs.

One significant limitation of our murine sarcoma models is their rapid tumor growth, which does not allow sufficient time for in vivo DAC+Enti treatment followed by immune priming before tumors have grown too large – not allowing time for the immune priming to generate effective immune responses. In vivo responses to these drugs have been more pronounced when multiple treatment cycles can be given, such as murine models of peritoneal ovarian carcinoma (18). We are working to establish additional sarcoma models that will permit longer in vivo treatment to better model this treatment strategy. In addition, while our studies define the impact of DAC+Enti treatment on sarcoma cells, we do not focus on how HMA and HDACi modulate other stromal and immune cells of the tumor microenvironment (TME). These drugs do induce important changes in the TME that will likely influence the effectiveness of such a strategy in patients (67). Other reports suggest beneficial effects of these drugs on the TME. Enti has been shown to inhibit the immunosuppressive function of myeloid-derived suppressor cells (MDSCs) in breast and pancreatic cancer (21). Low dose adjuvant epigenetic therapy using HMA and HDACi has also been shown to decrease trafficking of MDSCs to premetastatic niches in non-small cell lung cancer, decreasing lung metastases and improving overall survival (68). Clinical studies in sarcomas will also provide significant insights. We are currently analyzing the effects of HMA treatment in lung-relapsed osteosarcoma using samples from the recently completed trial of patients treated with neoadjuvant azacitidine and nivolumab (NCT03628209).

In summary, our study demonstrates that epigenetically regulated antigens can serve as potent T cell targets to increase the immunogenicity of poorly immunogenic sarcomas. We also provide evidence that mere expression of these antigens is not sufficient to generate immune activation, but that active immune stimuli such as a vaccine, or other locally delivered immune adjuvants, are required to generate T cell responses to these antigens. Finally, these epigenetically regulated antigens may be shared between different tumors and tumor types, broadening the applicability of this approach in sarcomas.

## Methods

### Sex as a biological variable

The murine tumor models used in our experiments originated from female mice. To not confound the effects of hypomethylating agents on sex chromosomes, the tumor challenge experiments were performed in female mice.

### Mouse and tumor models

C57BL/6 mice (obtained from Jackson Laboratories and bred in house) were housed in the animal vivarium at Johns Hopkins University and used according to the animal protocol approved by the Johns Hopkins University Animal Care and Use Committee. KP Sarc undifferentiated pleomorphic sarcoma cells were originally generated by injection of adenovirus expressing Cre-recombinase (Ad-Cre) into the muscle of *LSL-Kras*^G12D/+^*Trp53*^fl/fl^ mice fully back-crossed to a C57BL/6 genetic background (69) and generously provided by Dr. Jonathan Powell. Murine sarcoma M3-9-M cells were derived from a spontaneously arising sarcoma occurring in a C57BL/6 mouse transgenic for hepatocyte growth factor and heterozygous for mutated *p53* (37) and were generously provided by Dr. Crystal Mackall. KP Sarc and M3-9-M cells were cultured in RPMI 1640 medium (Invitrogen) supplemented with 10% heat-inactivated fetal bovine serum (FBS; Gemini Bio), antibiotics (penicillin G 100 units/mL, streptomycin sulfate 100 µg/mL, amphotericin B 0.25 µg/mL; Corning, 30-004-CL) and L-Glutamine (Corning, 25-005-CK) at 37 °C with 5% CO_2_.

KP Sarc and M3-9-M cells were treated *in vitro* with either 100 nM decitabine (DAC) (Selleckchem, S1200) for 72 h (drug added to culture every 24 h) or 2 µM entinostat (Enti) (Selleckchem, S1053) for 72 h drug exposure (drug added only on day 1). For the combination epigenetic treatment, cells were treated sequentially with DAC for 72 h followed by Enti for 72 h (DAC+Enti). KP Sarc cells were treated with 50 ng/mL murine IFN-γ for 24 h prior to MHC-I expression analysis and injection.

Tumors were established by orthotopic intramuscular injection of 2 x10^5^ tumor cells into the gastrocnemius. As tumors would grow from deep in the muscle, leg circumference was used to determine tumor size and was calculated by measuring 2 perpendicular axes of the tumor and using the formula for perimeter of an ellipse 2π√((major axis^2^+minor axis^2^)/2). Mice were euthanized when a maximum tumor diameter of 20 mm was reached or significant loss of mobility. Baseline leg circumference was about 17-19 mm. For tumor-free survival curves, a threshold of leg circumference >25 mm was used to declare the mouse tumor-bearing. Mice that rejected tumor formation were re-challenged with KP Sarc or M3-9-M cells 45 days post-initial challenge.

### Human sarcoma cell lines

Human sarcoma cell lines were acquired from the Childhood Cancer Repository (Lubbock, Texas, USA). CHLA-9 (RRID:CVCL_M150) was derived from *EWSR1::FLI1* translocation positive Ewing sarcoma at diagnosis (prior to treatment) (70). Rh30 (RRID:CVCL_0041) and Rh41 (RRID:CVCL_2176) are both *PAX3::FOXO1* translocation positive alveolar rhabdomyosarcoma derived respectively from bone marrow at diagnosis and from relapsed lung tumor. In vitro treatment with DAC and Enti was performed as described above.

### In vivo treatment

Each GVAX injection consists of 1×10^6^ KP Sarc or M3-9-M cells admixed with 1×10^6^ B78H1-GM-CSF secreting bystander cell line and irradiated with 50 Gy prior to injection. B78H1-GM-CSF line was titrated to secrete GM-CSF at 100 ng/1×10^6^ cells/24 h and were provided by Dr. Elizabeth Jaffee (36). Mice were given three GVAX injections subcutaneously in both hindlimbs and one forelimb. Cyclophosphamide was administered intraperitoneally (IP) as a 100 mg/kg dose 24 h prior to GVAX per standard protocol to provide transient regulatory T cell depletion (71). Anti-PD-1 (clone RMP1.14; BioXCell, BP0146) and anti-CTLA-4 (clone 9H10; BioXCell, BP0131) were given each at 200 µg per intraperitoneal injection, every three days until mice were sacrificed or tumors were cleared. T lymphocytes were depleted by administering anti-CD8^+^ (clone 2.43; BioXCell, BP0061) and anti-CD4^+^(clone GK1.5; BioXCell, BP0003-1) each at 200 µg per intraperitoneal injection twice per week. DAC (2.5 mg/kg) and Enti (5 mg/kg) were administered intraperitoneally and by oral gavage, respectively, daily for five days (21).

### Tumor Infiltrating Lymphocytes (TIL) Analysis

Tumors were harvested, weighed, finely minced, and digested for 30 minutes at 37 °C in 5 mL of digest media (RPMI with 5% heat inactivated FBS, 0.04 mg/mL DNAse I (Millipore Sigma, 10104159001), 225 units/mL collagenase I (Gibco, 17-018-029)) and passed through a 70 µM cell filter to obtain a single-cell suspension. TILs were enriched by overlay on Ficoll-Paque PLUS (Cytiva, 17544202) density gradient media. Cytokine production was induced by PMA/Ionomycin incubation for four h in the presence of monensin (BD Biosciences, 554724).

### Histone isolation and Acetylation analysis

Histones were isolated via acid extraction exactly as detailed previously (72). Briefly, cell pellets were washed and re-suspended in hypotonic buffer supplemented with protease inhibitor cocktails. Nuclear pellets were re-suspended in 0.2 N HCl and incubated overnight on a rotator. Histones were precipitated by adding trichloroacetic acid dropwise to a final concentration of 33%. Precipitated histones were washed with acetone, air-dried and were dissolved in 100 µL of water. Proteins were resolved on gradient gels (Bio-Rad) and analyzed as previously described (73). Acetylated histones were detected using Ac-H3 antibody (rabbit polyclonal; Millipore Sigma, 06-599). Total histones were detected using H3 (clone D1H2; Cell Signaling Technology, 4499S). Raw volumes of the bands were quantified on Bio-rad Chemi-Doc imaging system and were normalized to respective controls as described (74).

### Flow cytometry

Samples were analyzed on a BD FACSCelesta flow cytometer following staining. All samples were Fc blocked (BD Biosciences, clone 2.4G2, 553142) prior to staining and live/dead stained with eFluor780 Fixable Viability Dye (Invitrogen, 65-0865). Staining was done using the following fluorochrome-conjugated antibodies specific for: Foxp3 (PE; clone FJK-16s; Invitrogen, 12-5773-82), CD11b (PerCP-Cy5.5; clone M1/70; Invitrogen, 45-0112-82), CD4 (APC; clone RM4-5; BD Biosciences, 553051), CD45 (BV510; clone 30-F11; Biolegend, 563891), CD8α (BV605 or PerCP-Cy5.5; clone 53- 6.7; BD Biosciences, 563152 or 551162), IFN-γ (APC; clone XMG1.2; BD Bioscience, 554413), PD1 (BV605; clone J43; BD Biosciences, 563059), Tim3 (APC; clone RMT3-23; Biolegend, 119706), Lag3 (BV785; clone C9B7W; Biolegend, 125219), H-2 murine MHC I (PE; clone M1/42; Biolegend, 125506). For, intracellular staining we used eBioscience Transcription Factor Staining Buffer Set (Invitrogen, 00-5523-00) to stain Foxp3 and BD Cytofix/Cytoperm Fixation/Permeabilization Kit (BD Biosciences, 554714) to stain IFN-γ. Results were analyzed using FlowJo software (version 10.9.0).

### Immunohistochemistry

Formalin-fixed, paraffin-embedded (FFPE) tumor samples were sectioned into 4 µm tissues mounted on charged Superfrost slides (Fisher Scientific, 22-037-246). Antigen retrieval was done using 1X DAKO Retrieval buffer pH6 (DAKO, S1699). We incubated the samples with the primary rabbit polyclonal anti-Dazl antibody (Abcam, ab34139) for 45 minutes, at a concentration of 1 µg/mL. The secondary antibody was incubated for 30 minutes. DAB (DAKO, SK-4105) was applied for 45 seconds. Slides were counterstained with 1:5 hematoxylin (VWR, 95057-844). Slides were scanned digitally by Scanscope XT and analyzed using Indica Labs HALO pathology software. We manually delineated tumor borders and used HALO to determine Dazl-positive cell density per tumor area.

### RNA isolation and Real Time RT-PCR

Total RNA was isolated using a RNeasy Mini kit (Qiagen, 74104) as described (75). Purified RNA was converted to cDNA using the SuperScript III Reverse Transcriptase (Fisher Scientific, 18-080-093). Quantitative PCR (qPCR) was performed by standard techniques with specified Taqman primer/probe sets for Dazl (Assay ID: Mm01273546_m1, ThermoFisher Scientific) and Trap1a (Assay ID: Mm00495785_m1, ThermoFisher Scientific). We quantified relative gene expression levels standardized to eukaryotic 18S rRNA endogenous control (Applied Biosystems, ThermoFisher Scientific, 4319413E) in each sample using the ΔΔCT method (76).

### Bulk RNA sequencing (RNA-seq)

RNA-seq library preparation, sequencing, and alignment were performed as previously described (77) using the NEBNext Ultra II Directional RNA Library Prep Kit and Illumina HiSeq 4000 platform. Differential gene expression analysis was performed using R version 4.4.2 (2024-10-31) and the DESeq2 package (1.46.0) on a summarized experiment object derived from STAR-aligned BAM files. Raw counts were normalized using DESeq2, and a regularized log (rlog) transformation was applied to stabilize variance across samples. Volcano plots were generated for each condition: DAC, Enti, and DAC+Enti, compared to untreated (CTL) KP Sarc using ggplot2 in R, applying thresholds of adjusted p-value (padj) < 0.05 and log₂ fold change (log₂FC) < -2.5 or > 2.5 to define significantly differentially expressed genes (DEGs). The same analysis was done for DAC+Enti-treated human sarcoma cell lines, RH30, RH41, and CHLA-9. To delineate subsets of upregulated genes, we categorized genes based on their expression in the control condition: those with average normalized counts below 50 in CTL but above 50 in DAC+Enti were considered “low in CTL” and those with counts above 50 in CTL were “high in CTL.” A heatmap of all significantly upregulated genes low in CTL was generated across all conditions using the pheatmap R package (1.0.12). The heatmap was row-scaled (z-score normalized), and hierarchical clustering was applied to both rows and columns using Euclidean distance and complete linkage. A blue-to-red color gradient was used to represent low to high expression values. Gene set enrichment analysis was performed using the gseGO function from the clusterProfiler R package (4.14.6) on the DESeq2-derived gene-level statistics (Wald test t-values). Enrichment was tested against the Gene Ontology (GO) Biological Process (BP) database and assessed using normalized enrichment scores (NES) and padj. Significantly enriched pathways were visualized using a custom ggplot2-based bubble plot, where bubble size reflected the NES and color scale denoted -log_10_(padj). Principal component analysis (PCA) was performed to assess global transcriptomic changes of RH30, RH41, and CHLA-9 based on DESeq2 expression counts of DEGs selected for padj below 0.05. Genes shared across cell lines were scaled using the prcomp function in R.

### Whole-genome bisulfite sequencing

Whole-genome bisulfite sequencing (WGBS) libraries were prepared from control and treated tumor cells as previously described (78) using the NEBNext Ultra DNA Library Prep kit. Transcription start sites (TSS) were defined using the TxDb.Mmusculus.UCSC.mm9 gene annotation in R. Mean methylation levels (MML) around TSS were determined using informME (79).

### Statistics

All dot plots show mean ± SD. One-way ANOVA with Tukey’s multiple comparison test and two-sided log-rank test were used. A p-value < 0.05 was considered statistically significant. Statistical analyses were performed using GraphPad Prism 10. Other statistical methods were used as described elsewhere.

### Study approval

Animal studies were conducted according to approved protocols by the Johns Hopkins University Animal Care and Use Committee. Appropriate biosafety approvals for laboratory work and reagents were maintained and renewed annually with the Johns Hopkins University Biosafety Office.

## Supporting information

Supporting Data values

## Data availability

Raw data for each figure and survival data are provided in the supplemental file “Supporting Data Values.” RNA-seq expression data and whole-genome bisulfite sequencing data from treated tumor cell lines will be deposited in the Gene Expression Omnibus (GEO) data repository before revision.

## Author Contributions

Designed research studies: AR, HRG, EER, MAK, BHL. Conducted experiments: AR, HRG, EER, ANML, MJP, SG, MD, BHL. Acquired data: AR, HRG, EER, ANML, MJP, SG, MD, MIB, MAK. Analyzed data: AR, HRG, EER, ANML, MJP, SG, MD, MIB, MAK. Wrote the manuscript: AR, HRG, EER, MAK, BHL. Manuscript editing: AR, HRG, EER, ANML, MJP, SG, MD, NJL, MAK, BHL. Obtained grant funding: NJL, BHL. Co-first authors AR and HRG – HRG contributed more to data generation, AR contributed data and most to data analysis, figure generation, and manuscript completion with author order based on most recent contributions and completion of the manuscript for publication.

## Acknowledgements

none

---

**Supplemental Figure 1:**
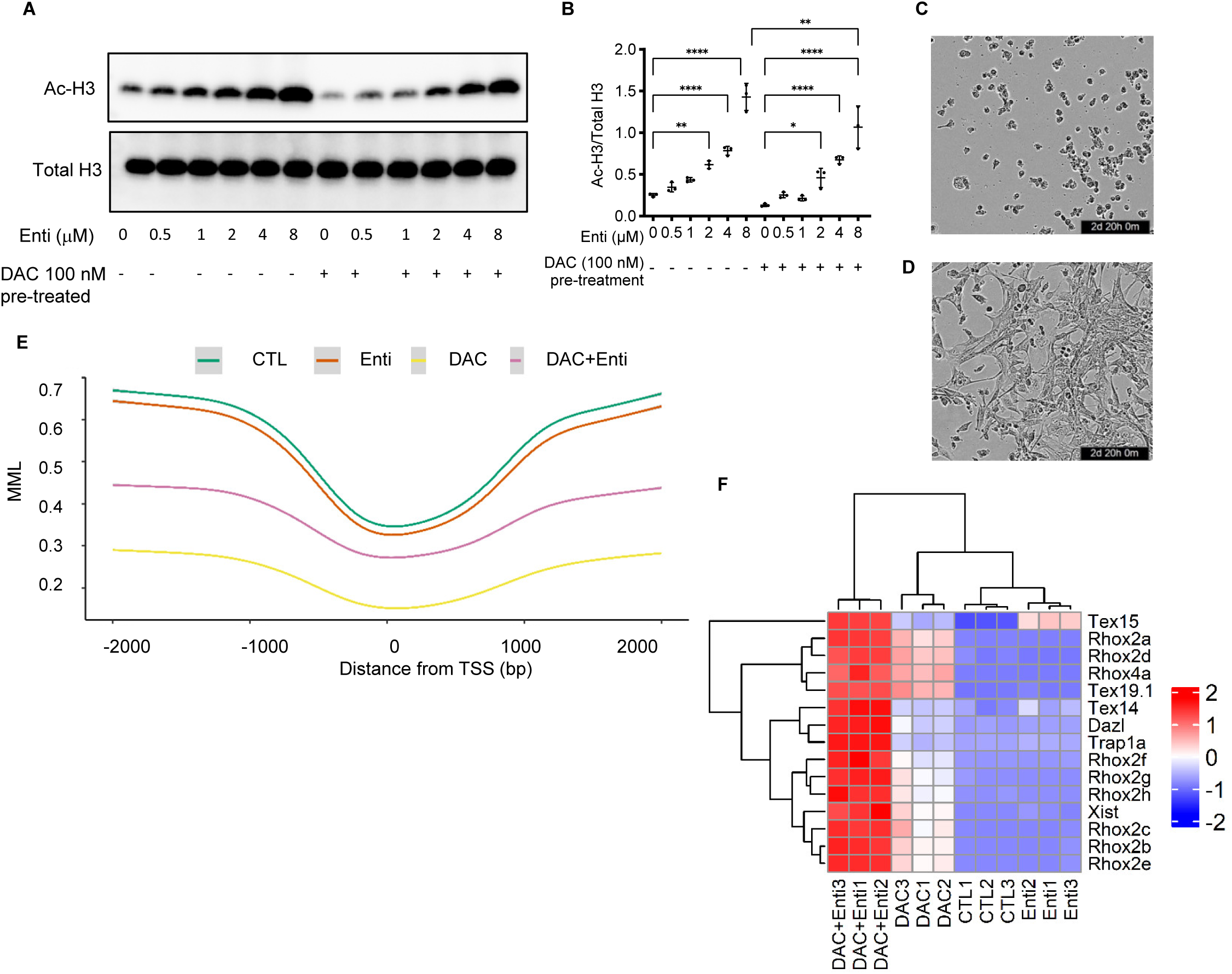
Sequential epigenetic therapy induces global histone, genome, gene expression, and morphologic changes in KP Sarc cells. *(A)* Western blot of Ac-H3 and total H3 in KP Sarc cells incubated with increasing concentrations of Enti for 72 h, with or without pre-treatment with DAC 100 nM for 72 h. (*B)* Densitometric analysis of Ac-H3 normalized to total H3 from *(A)* with results combined from three experimental replicates. Comparisons were made between the various Enti concentrations within each of the DAC-pretreatment groups; as well as between the same Enti concentration within the two groups. Only significant comparisons are shown. The values represent the mean ± SD following one-way ANOVA, * p< 0.05, ** p< 0.001, *** p< 0.0005, **** p< 0.0001. Representative light microscopy image of KP Sarc cells in culture *(C)* and changes to KP Sarc cell morphology after incubation with Enti *(D)* for 72 h. *(E)* DNA mean methylation levels (MML) of control (CTL), DAC, Enti, or DAC+Enti treated KP Sarc as analyzed within 2000 kb of ref-seq transcription start sites (TSS). *(F)* Heatmap of the cancer testis antigens (CTAs) from the upregulated, low in CTL genes identified in Figure 1C. Count data were normalized using DESeq2’s regularized log transformation (rlog) and scaled using Z-score. Hierarchical clustering was performed using Euclidean distance and complete linkage within the R pheatmap package.

**Supplemental Figure 2:**
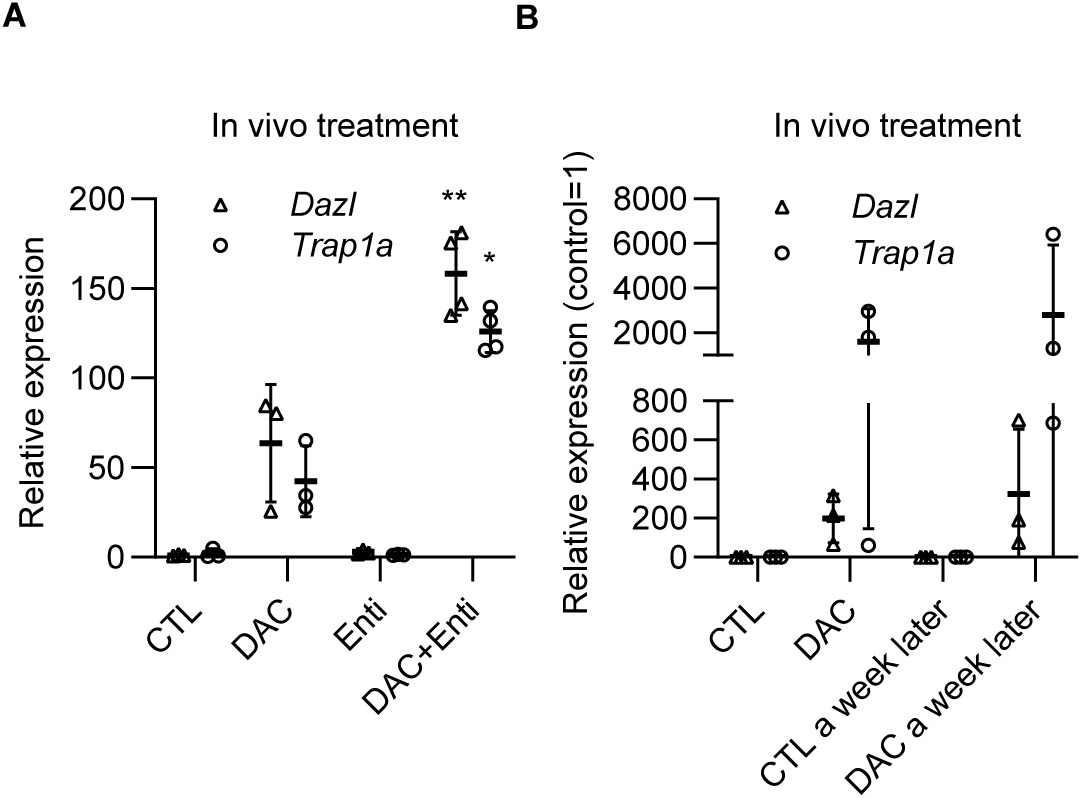
qPCR analysis of mRNA levels of CTAs *Dazl* and *Trap1a* in KP Sarc tumors treated in vivo with *(A)* either DAC (2.5 mg/kg/dose) intraperitoneally x 5 days or Enti (5 mg/kg/dose) by oral gavage x 5 days or sequentially DAC+Enti or *(B)* only with DAC and harvested at end of treatment or a week later. The values represent the mean ± SD as determined by Kruskal-Wallis test with *p<0.05, **p<0.01 as compared to vehicle control.

**Supplemental Figure 3:**
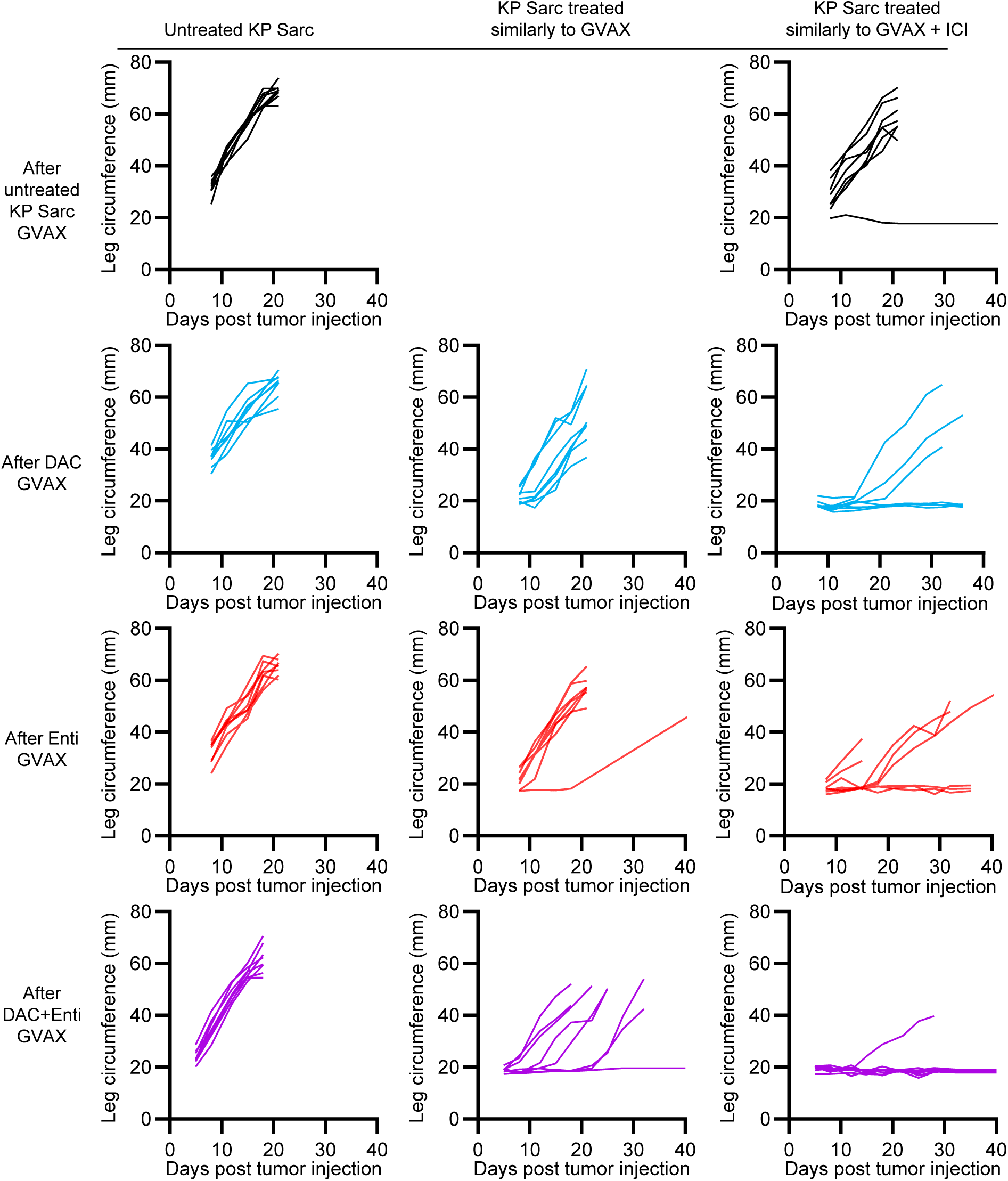
Spider plots showing individual mice KP Sarc growth after GVAX injection.

**Supplemental Figure 4:**
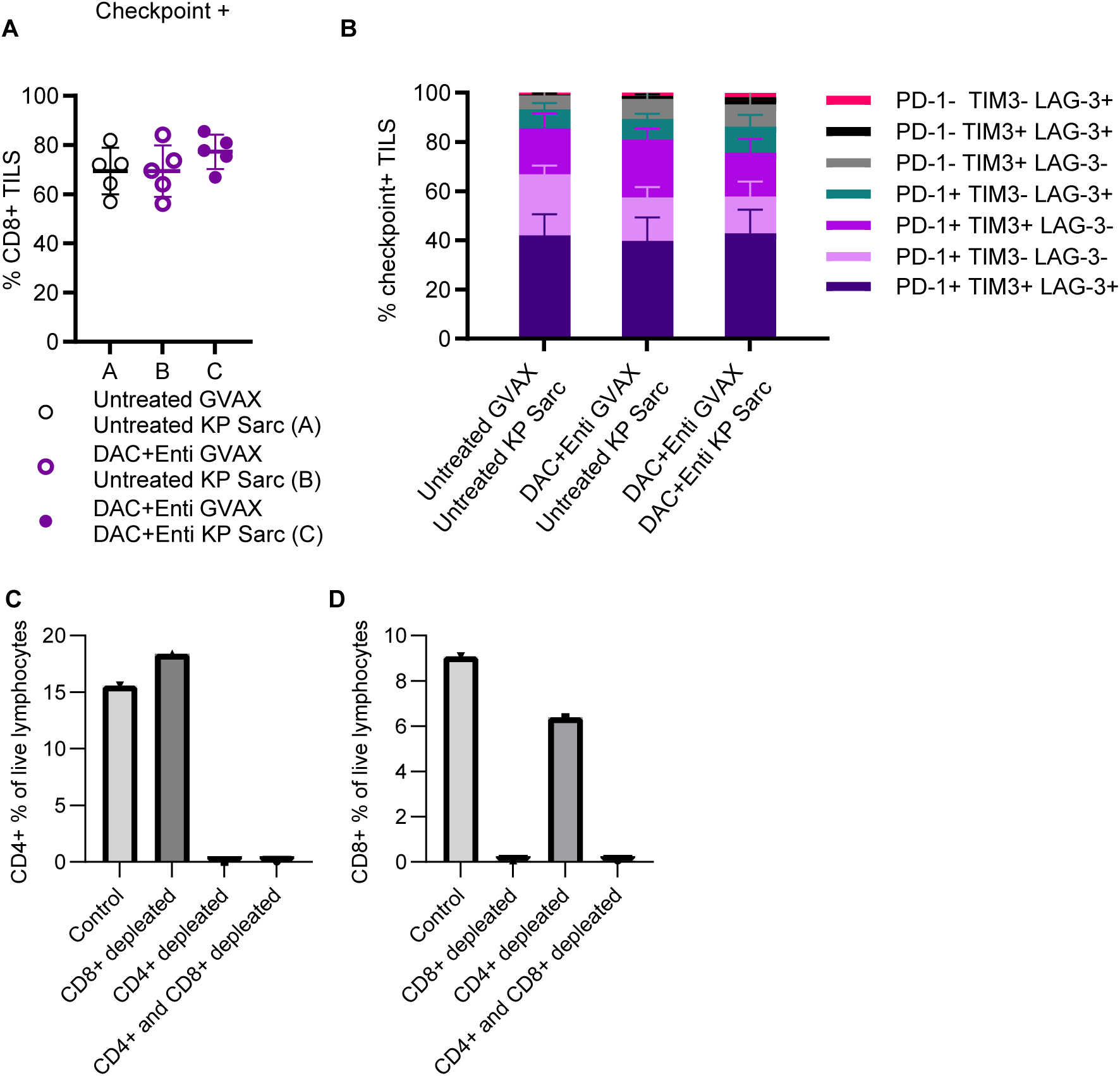
Quantification using flow cytometry of checkpoint^+^ T cells *(A)* and the stacked bar figures showing the relative composition of cells expressing various combinations of PD-1, Tim3 and Lag3 *(B)*. The values represent the mean +/- SD. *p<0.05 as determined by one-way ANOVA with Tukey’s multiple comparison test. Quantification of CD4^+^ *(C),* and CD8^+^ T cells *(D),* in splenocytes isolated from mice which received CD4^+^ and/or CD8^+^ depleting antibodies.

**Supplemental Figure 5:**
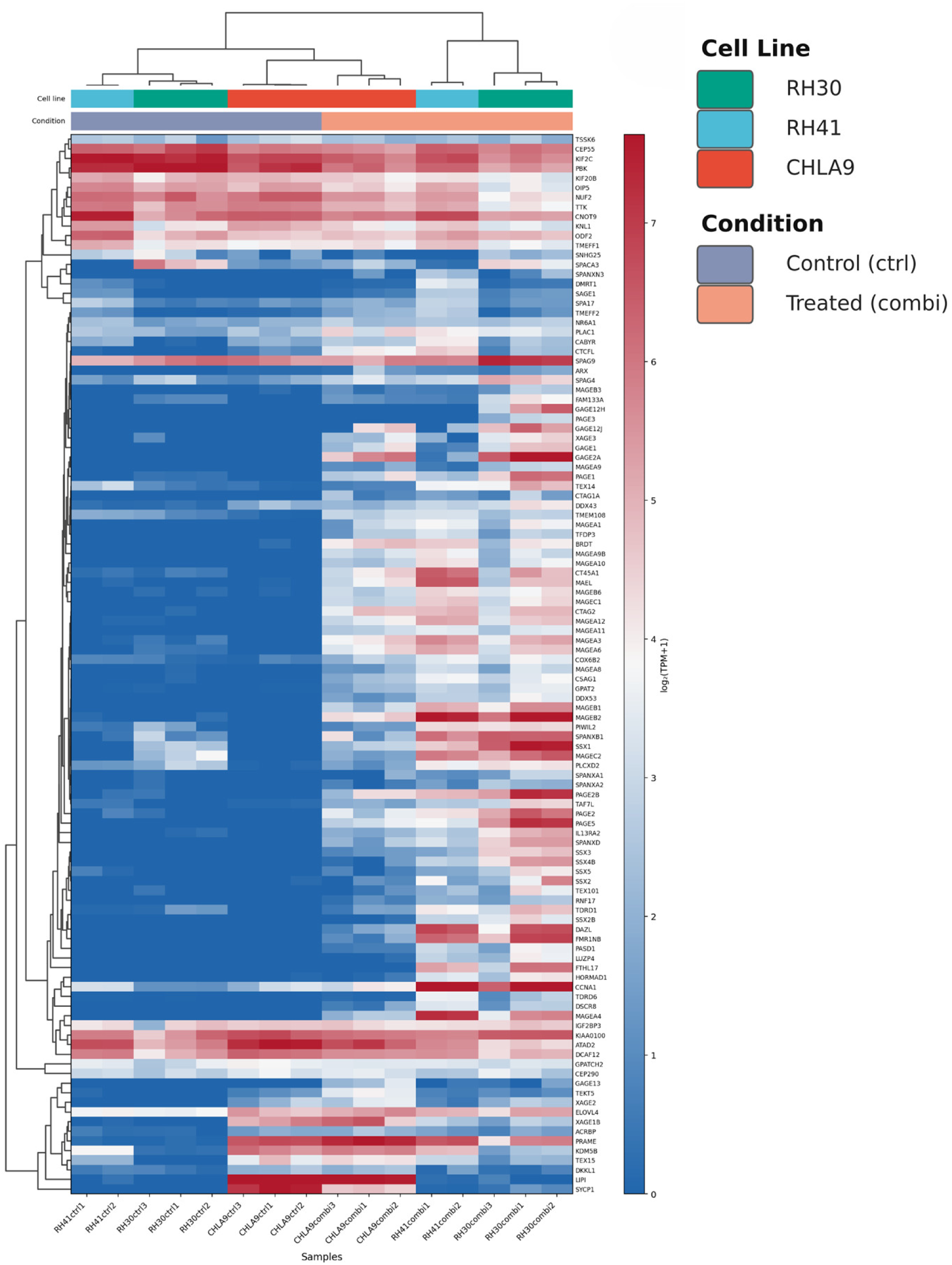
Heatmap displaying normalized gene expression data (log_2_(TPM+1)) of defined Cancer Testes Antigens (CTAs) (http://www.cta.lncc.br/list_of_genes.html) in DAC+Enti (combi) vs control (ctrl) untreated RH30, RH41, and CHLA-9 cells. Gene list limited to CTAs with TPM >5 in at least 1 sample. Clustering by average linkage with correlation distance, applied independently to both genes (rows) and samples (columns).

